# Cas9/sgRNA-mediated genome editing of citrus via mature tissue transformation enables both high-efficacy genome editing and early flowering

**DOI:** 10.64898/2026.04.20.719603

**Authors:** Hongge Jia, Zhuyuan Hu, Hao Wu, Yucheng Duan, Janice Zale, Nian Wang

**Affiliations:** Citrus Research and Education Center, Institute of Food and Agricultural Sciences, University of Florida, Lake Alfred, FL. U.S.A.; Department of Microbiology and Cell Science, Institute of Food and Agricultural Sciences, University of Florida, Lake Alfred, FL. U.S.A.; Plant Molecular and Cellular Biology Graduate Program, Institute of Food and Agricultural Sciences, University of Florida, Lake Alfred, FL. U.S.A.

**Keywords:** mature tissue transformation, citrus, CRISPR, genome editing, juvenility, flowering

## Abstract

CRISPR genome editing has shown tremendous potential in genetic improvement of citrus. So far, citrus genome editing has been conducted using juvenile tissues resulting in genome-edited citrus plants that require multiple years before they can produce flowers and fruit. Here we tested whether citrus genome editing via mature tissue transformation can overcome such a hurdle. *CsLOB1* is a susceptibility gene for citrus canker caused by *Xanthomonas citri* subsp. *citri* (Xcc). The transcription activator-like effector PthA4 of Xcc activates *CsLOB1* by binding to the effector-binding element in its promoter (EBE_pthA4_-CsLOBP). In Valencia sweet orange, two *CsLOB1* promoter alleles are present: TI CsLOBP, and TII CsLOBP. We specifically utilized a CRISPR/Cas9 construct (GFP-p1380N-Cas9/sgRNA:CsLOBP2) targeting EBE_pthA4_ in TI CsLOBP but not TII CsLOBP to test genome editing efficacy and off-target mutations. GFP-p1380N-Cas9/sgRNA:CsLOBP2 function was first validated using Xcc-facilitated agroinfiltration in Valencia leaves. The construct was subsequently introduced into Valencia mature internodal stem segments via *Agrobacterium*-mediated transformation, generating three independent transgenic lines (#V2, #V3 and #V5). Targeted mutations in EBE_pthA4_-TI CsLOBP were detected in all three lines with mutation frequencies of 100%, 21.43% and 41.94% in #V2, #V3 and #V5, respectively, while no mutations were detected in TII CsLOBP. Infection with XccΔpthA4:dCsLOB1.3, carrying a designer TALE that specifically activates TI CsLOBP, resulted in reduced canker symptoms in #V2. Importantly, all three EBE_pthA4_-TI CsLOBP edited lines flowered within 15 months. In sum, these results demonstrate that CRISPR/Cas9-mediated genome modification through mature citrus transformation can achieve high genome editing efficacy and overcome the juvenility.

## Introduction

Citrus fruits are rich in nutrients, including vitamins, fiber, and sugars, making them among the most widely consumed fruits worldwide. However, global citrus production is severely threatened by many devastating diseases including Huanglongbing (HLB) and environmental stresses. Therefore, genetic improvement of citrus is urgently needed to overcome those challenges. CRISPR-Cas-mediated genome editing has emerged as one of the most promising approaches for breeding disease-resistant citrus varieties. Compared with traditional hybrid breeding, CRISPR-Cas-based breeding enables faster and more precise genetic improvement with predictable outcomes (Molla et al., 2021).

CRISPR-Cas genome editing was first used to mutate the *CsPDS* gene in citrus (Jia and Wang, 2014). Subsequently, the PthA4 effector binding elements (EBE_pthA4_) region in the promoter and the coding sequence of *CsLOB1*, the citrus canker susceptibility gene, were modified to increase disease resistance to citrus canker caused by *Xanthomonas citri* subsp. citri (Xcc) (Jia et al., 2017b; Peng et al., 2017). PthA4 is the most important virulence factor of Xcc, which activates *CsLOB1* by binding to EBE_pthA4_ to induce disease symptoms (Hu et al., 2014). To date, several genome-editing platforms have been applied to citrus genetic improvement and functional studies, including base editors derived from SpCas9/gRNA, SpCas9/gRNA from *Streptococcus pyogenes*, SaCas9/gRNA from *Staphylococcus aureus*, and LbCas12a/crRNA from *Lachnospiraceae* bacterium in previous work (Huang et al., 2021; Huang et al., 2022; Huang et al., 2020; Jia et al., 2019; Jia and Wang, 2014; Jia et al., 2017a; LeBlanc et al., 2018; Prado et al., 2024). However, most previous studies relied on immature citrus epicotyls or callus tissues as explants for transformation. A major limitation of regenerants derived from immature tissues is the prolonged juvenile phase. Typically, citrus trees grown from juvenile tissues take 5 to 10 years to transit from the juvenile phase to maturity, which is necessary for flowering and fruiting (Gmitter et al., 2012). This extended juvenile period substantially delays the evaluation and deployment of new citrus cultivars generated through CRISPR-Cas-mediated breeding.

To circumvent the long juvenile phase, mature tissue transformation was developed for citrus genetic engineering in 1998 (Cervera et al., 1998). Mature citrus transformation involves the genetic modification of mature tissues rather than juvenile tissues (Cervera et al., 1998; Orbović et al., 2015). In this approach, clean apical tip meristems from mature trees are grafted onto rootstocks maintained under sanitary conditions to generate rejuvenated shoots for transformation. Here, mature internodal stem segments from Valencia sweet orange were used as explants to evaluate the efficiency and off-target mutations of Cas9/sgRNA-mediated genome editing of EBE_pthA4_-TI CsLOBP in mature citrus tissues.

## Method

### Plasmid construction

The CaMV 35S promoter was amplified with primers CaMV35-5-*Xho*I (5′-A<u>CTCGAG</u>ACTAGTACCATGGTGGACTCCTCTTAA-3′) and sgRNA-CsLOBP2-1 (5′-phosphorylated-GAACTTTGTTCCTCTCCAAATGAAATGAACTTC-3′), while the sgRNA-NosT fragment was amplified using sgRNA-CsLOBP2-2 (5′-phosphorylated-AAGGCAAAAGGTTTTAGAGCTAGAAATAGCAA-3′) and NosT-3-*Asc*I (5′-ACCTGGGCCC<u>GGCGCGCC</u>GATCTAGTAACATAGATGA-3′). The resulting fragments were assembled via three-way ligation: *Xho*I-digested CaMV35S and *Asc*I-cut sgRNA-NosT were inserted into *Xho*I-*Asc*I-treated GFP-p1380N-Cas9/sgRNA:cslob1 to generate GFP-p1380N-Cas9/sgRNA:CsLOBP2. The GFP-p1380N-Cas9/sgRNA:cslob1 vector was described previously (Jia et al., 2017), and the binary vector p1380-AtHSP70BP-GUSin was developed earlier (Jia and Wang, 2014b).

Binary vectors were introduced into *Agrobacterium tumefaciens* strain EHA105 by the freeze-thaw method, and recombinant cells were subsequently used for Xcc-facilitated agroinfiltration or mature citrus transformation.

### Xcc-facilitated agroinfiltration in Valencia

Valencia plants were grown in a greenhouse at 25-30□°C and pruned to ensure uniform growth prior to Xcc-facilitated agroinfiltration. Agroinfiltration was performed as previously described (Jia and Wang, 2014b). Leaves were first pre-treated with actively growing Xcc resuspended in sterile tap water (5×10^8^ CFU/ml). After 24□h, *Agrobacterium tumefaciens* cells carrying GFP-p1380N-Cas9/sgRNA:CsLOBP2 or p1380-AtHSP70BP-GUSin were infiltrated into the same Xcc-pretreated areas. Four days post-infiltration, leaves were harvested for GFP detection or genomic DNA extraction.

### *Agrobacterium*-mediate mature Valencia transformation

Mature citrus transformation was carried out as described previously (Wu et al., 2015). Mature *Valencia* internodal stem segments were used as explants and transformed with *Agrobacterium tumefaciens* carrying the binary vector GFP-p1380N-Cas9/sgRNA:CsLOBP2. Transgenic plants were screened for GFP fluorescence, and GFP-positive lines were further confirmed by PCR using primers Npt-5 (5′-ATTGAACAAGATGGATTGCACG-3′) and 35T-3 (5′-TTCGGGGGATCTGGATTTTAGTAC-3′).

### Valencia CsLOBP sequencing and indel analysis

Using a CTAB-based protocol (Huang et al., 2020), genomic DNA was extracted from wild type Valencia, or transgenic Valencia plants (#V2, # V3, # V5), or the Valencia leaves treated by Xcc-facilitated agroinfiltration with GFP-p1380N-Cas9/sgRNA:CsLOBP2-transformed *Agrobacterium* (Fig. 1c). Using 100 ng of genomic DNA as template, PCR was used to amply Valencia CsLOBP with the Q5 Hi Fidelity DNA polymerase (New England Biolabs) and a pair of primers, LOBP3 (5′-AGGTAAGCTTATTCATATTAACGTTATCAATGATT-3′) and LOBP2 (5′-ACCT<u>GGATCC</u>TTTTGAGAGAAGAAAACTGTTGGGT-3). PCR products were cloned via blunt-end ligation into the PCR-BluntII-TOPO vector (Life Technologies) and sequenced. Sequencing results were analyzed using Chromas Lite, and indels were identified by manual inspection.

**Figure 1.**
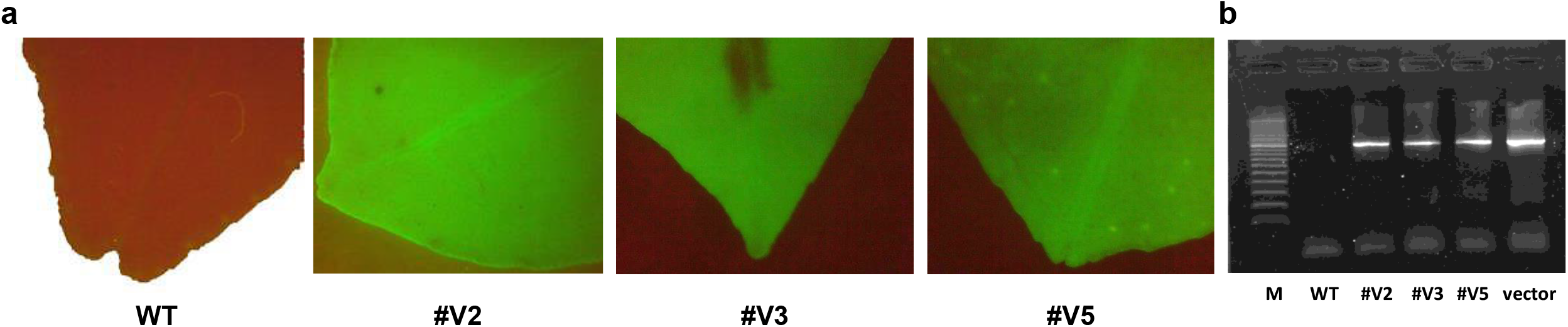
GFP detection and PCR confirmation of GFP-p1380N-Cas9/sgRNA:CsLOBP2-transgenic Valencia plants generated via mature tissue transformation. (a) GFP fluorescence in Valencia plants transformed with GFP-p1380N-Cas9/sgRNA:CsLOBP2. Three independent transgenic lines (#V2, #V3, and #V5) exhibited GFP-positive signals, whereas wild-type Valencia served as a GFP-negative control. (b) PCR analysis of transgenic Valencia lines using primers Npt-5 and 35T-3. A 980-bp amplicon was detected in transgenic plants and the plasmid control (GFP-p1380N-Cas9/sgRNA:CsLOBP2), but not in wild type Valencia sweet orange. M, DNA ladder; WT, wild type.

### Canker symptom assay in citrus

Valencia sweet orange (*Citrus sinensis*) and transgenic Valencia lines were grown in a glasshouse and pruned prior to pathogen inoculation. Leaves of the same developmental stage were inoculated with Xcc or XccΔpthA4:dCsLOB1.3 resuspended in sterile tap water (5×10^8^ CFU/ml). Canker symptoms were monitored at different time points, and representative photographs were recorded.

### GFP detection

GFP fluorescence was visualized using an Omax camera mounted on a Zeiss Stemi SV11 dissecting microscope. Leaves of Valencia treated with Xcc-facilitated agroinfiltration or transformed with GFP-p1380N-Cas9/sgRNA:CsLOBP2 were illuminated using a NIGHTSEA Stereo Microscope Fluorescence Adapter. Fluorescence images were captured with Omax ToupView software.

## Results

Based on the genealogy of the citrus family (Wu et al., 2018), two types of *CsLOB1* promoter (Type I CsLOBP and Type II CsLOBP) are expected to exist in Valencia sweet orange. PCR amplification and Sanger sequencing of the CsLOBP from Valencia indeed showed two different types. Among 22 randomly selected colonies, fourteen contained the mandarin-derived CsLOBP, designated as TI CsLOBP, and eight colonies contained the pummelo-derived CsLOBP, designated as TII CsLOBP (Supplementary Fig. 1a). There is an additional nucleotide (C) located immediately downstream of the EBE_pthA4_ in the TI CsLOBP (Supplementary Fig. 1a). Consistently, two types of CsLOBP are also present in Duncan grapefruit, a hybrid cultivar derived from Valencia sweet orange and pummelo.

To test the genome editing efficacy via mature tissue transformation, we used the vector GFP-p1380N-Cas9/sgRNA:CsLOBP2 to specifically target EBE_pthA4_-TI CsLOBP but not EBE_pthA4_-TII CsLOBP (Supplementary Figs. 1b and 1c) (Jia et al., 2016). This design enabled us to test the genome editing efficacy and off-target mutations against EBE_pthA4_-TII CsLOBP. The sgRNA in GFP-p1380N-Cas9/sgRNA:CsLOBP2 targets a 20-nt sequence within EBE_pthA4_-TI CsLOBP, located 2 bp closer to the TATA box than the target site of p1380N-Cas9/sgRNA:CsLOBP1 (Jia et al., 2016). To verify its genome editing efficacy, Xcc-facilitated agroinfiltration of Valencia leaves was initially employed. A total of 74 colonies were randomly selected for sequencing. Among 43 TI CsLOBP clones, three contained Cas9/sgRNA:CsLOBP2-mediated indels, whereas no modifications were detected among 31 TII CsLOBP clones (Supplementary Fig. 2). These results indicate that GFP-p1380N-Cas9/sgRNA:CsLOBP2 specifically targets TI CsLOBP.

Next, Valencia mature internodal stem segments were subjected to *Agrobacterium*-mediated transformation as previously described (Wu et al., 2015). Three GFP-positive lines (#V2, #V3, and #V5) were successfully generated (Fig. 1a). PCR analysis confirmed the presence of the transgene in regenerated Valencia transgenic lines (Fig. 1b). Sanger sequencing was performed to analyze Cas9/sgRNA:CsLOBP2-induced indels in lines #V2, #V3, and #V5. For each line, 50 colonies were randomly selected for sequencing. For #V2, 21 mutated EBE_pthA4_-TI CsLOBP and 29 WT EBE_pthA4_-TII CsLOBP were detected among 50 colonies (Fig. 2). For #V3, 6 mutated EBE_pthA4_-TI CsLOBP, 22 WT EBE_pthA4_-TI CsLOBP and 22 WT EBE_pthA4_-TII CsLOBP were detected (Fig.3). For #V5, 13 mutated EBE_pthA4_-TI CsLOBP, 18 WT EBE_pthA4_-TI CsLOBP and 19 WT EBE_pthA4_-TII CsLOBP were detected (Fig. 4). These results were consistent with the results obtained from Xcc-facilitated agroinfiltration (Supplementary Fig. 2). Accordingly, the Cas9/sgRNA:CsLOBP2-mediated mutation rates of the Type I allele were 100% (#V2), 21.43% (#V3), and 41.94% (#V5), and no off-target mutations were detected for EBE_pthA4_-TII CsLOBP.

**Figure 2.**
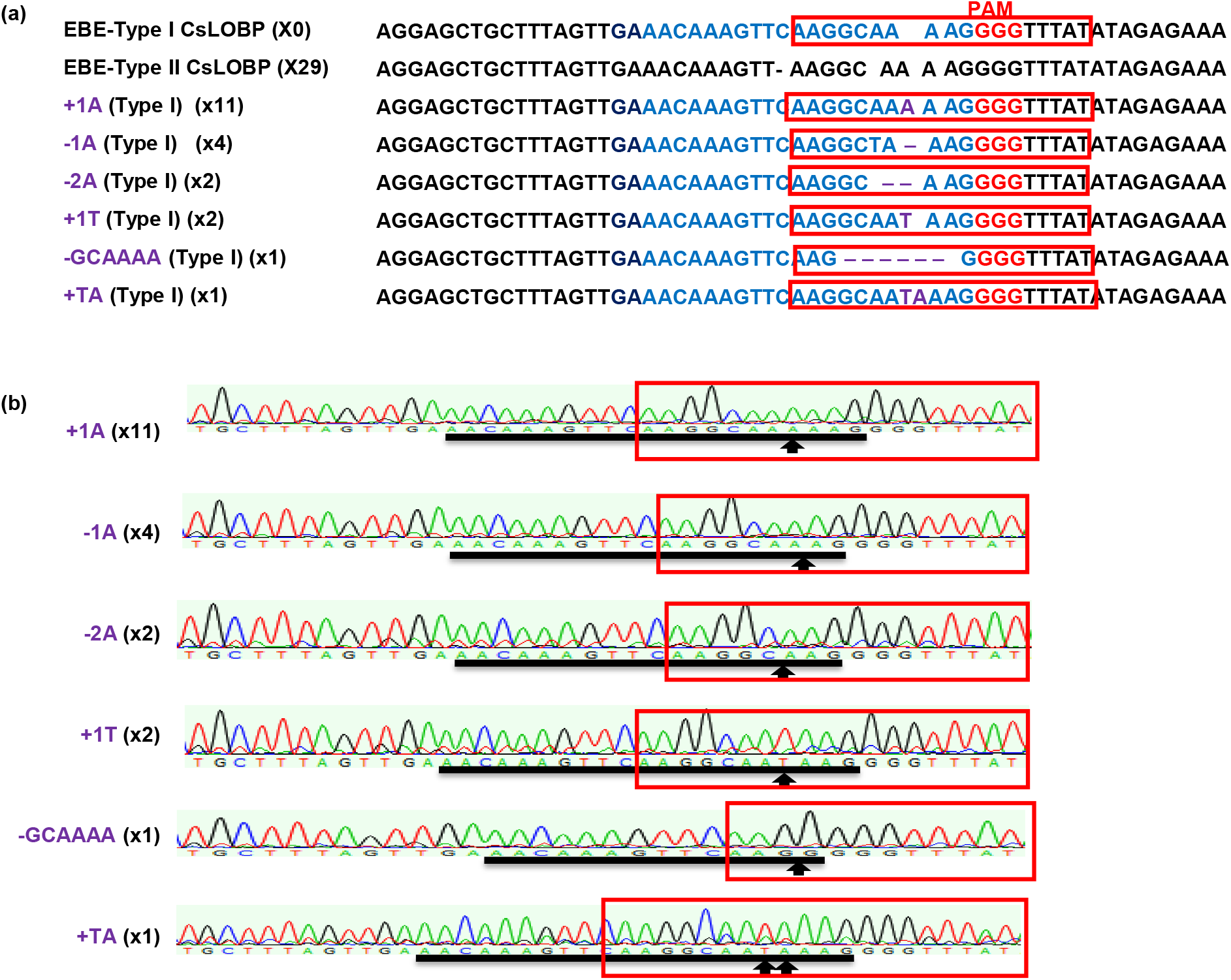
Cas9/sgRNA:CsLOBP2-mediated modification of EBE_PthA4_-TI CsLOBP in transgenic Valencia line #V2 generated via mature tissue transformation. (a) Indels induced by Cas9/sgRNA:CsLOBP2 in the EBE_PthA4_-TI CsLOBP in transgenic Valencia line #V2. The sgRNA target sequence was highlighted in blue, and mutations are indicated in purple. The protospacer-adjacent motif (PAM) was indicated in red. The PthA4 effector binding elements were marked with red rectangles. Among 50 sequenced colonies, 29 correspond to Type II CsLOBP and 21 to mutant Type I CsLOBP. (b) Representative chromatograms of Type I CsLOBP and its edited variants. The sgRNA target region was indicated by a black line, and mutation sites were marked by arrows.

**Figure 3.**
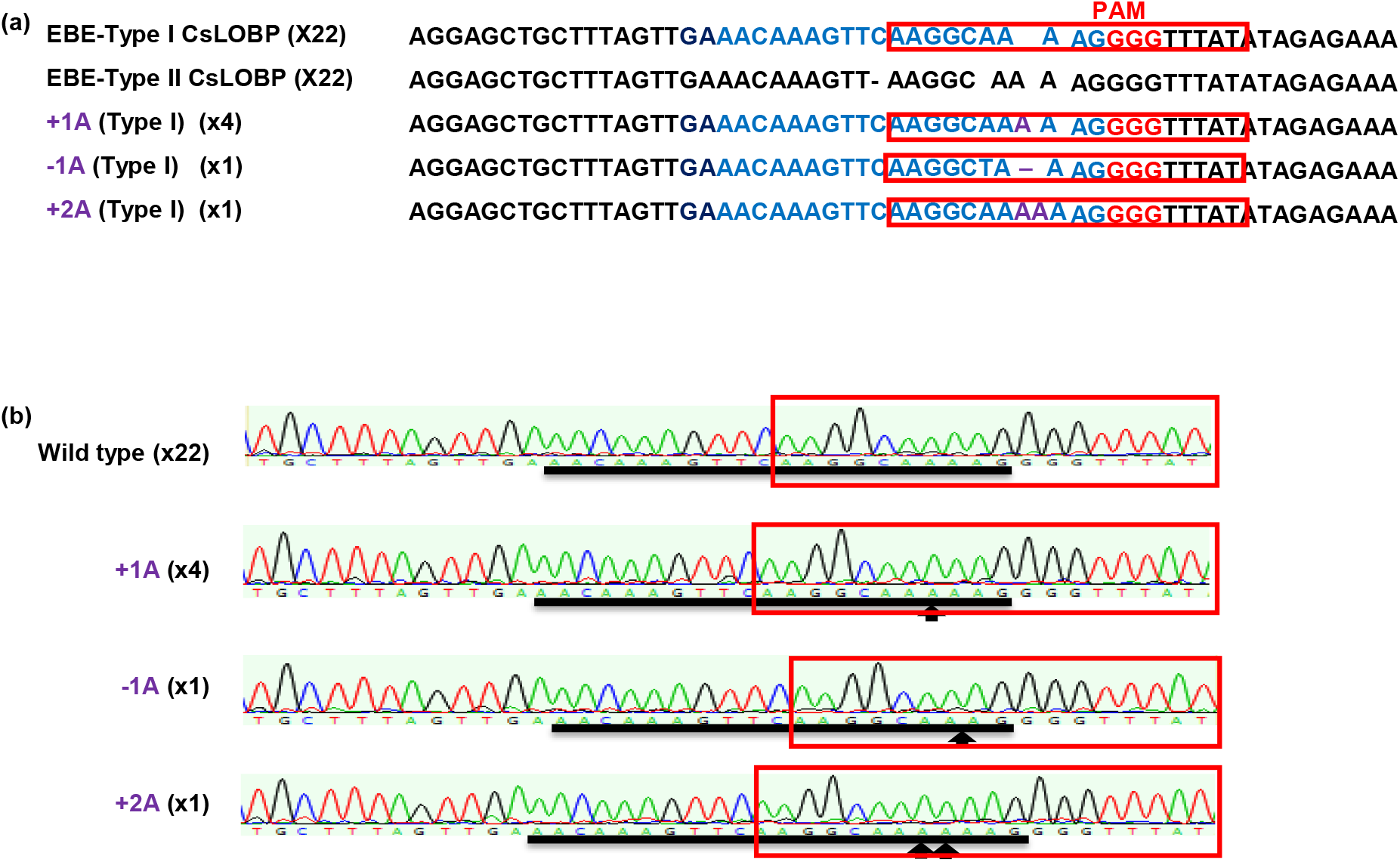
Cas9/sgRNA:CsLOBP2-directed modification of EBE_PthA4_-TI CsLOBP in transgenic Valencia line #V3 generated via mature tissue transformation. (a) Indels induced by Cas9/sgRNA:CsLOBP2 in the EBE_PthA4_-TI CsLOBP in transgenic Valencia line #V3. The sgRNA target sequence was highlighted in blue, and mutations are indicated in purple. The protospacer-adjacent motif (PAM) was indicated in red. The PthA4 effector binding elements were marked with red rectangles. Among 50 colonies sequenced, there were 22 Type I CsLOBP, 22 Type II CsLOBP, and 6 mutant Type I CsLOBP. (b) Representative chromatograms of Type I CsLOBP and its edited variants. The sgRNA target region was indicated by a black line, and the mutant sites were highlighted by arrows.

**Figure 4.**
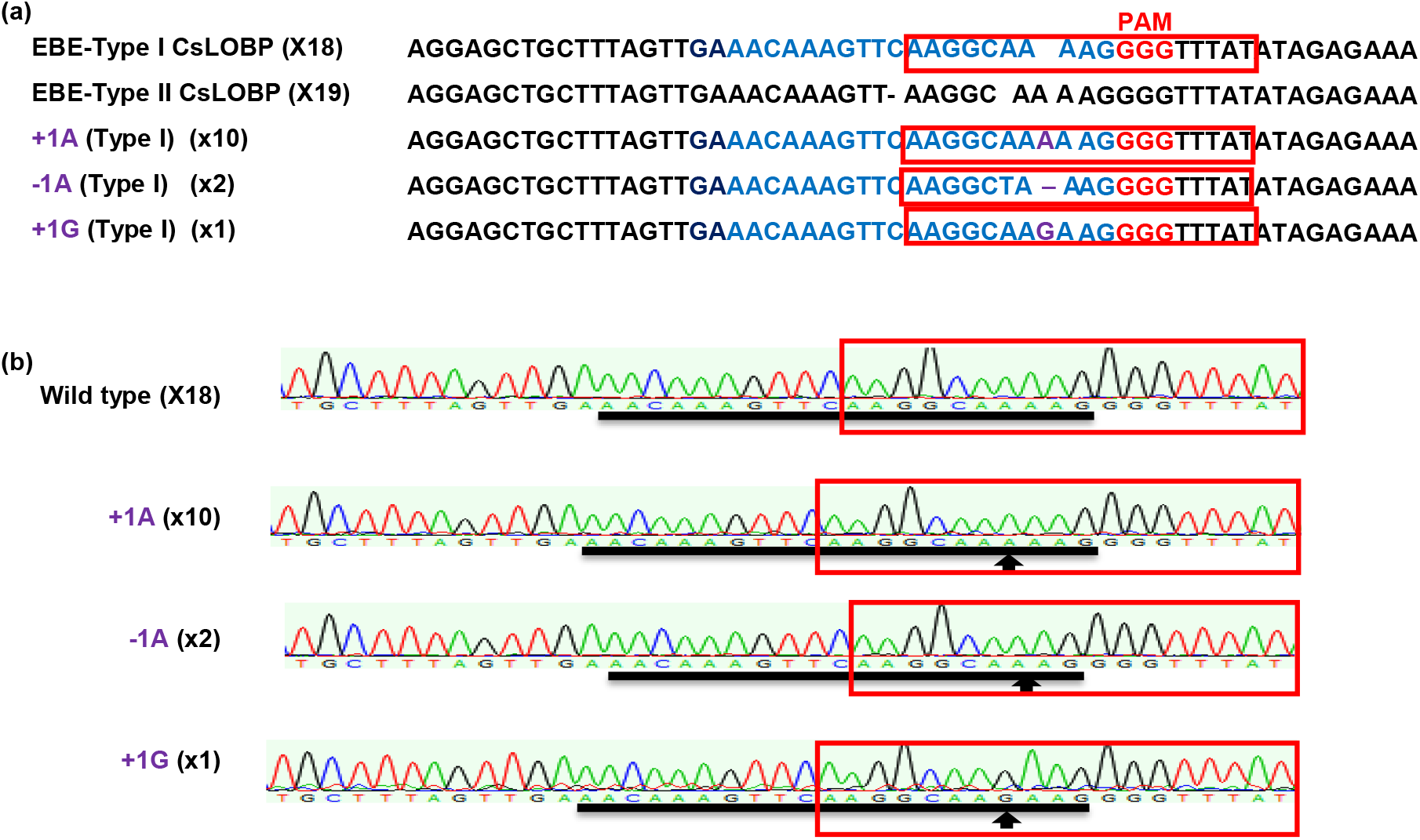
Cas9/sgRNA:CsLOBP2-mediated modification of EBE_PthA4_-TI CsLOBP in transgenic Valencia Line #V5 generated via mature tissue transformation. (a) Cas9/sgRNA:CsLOBP2-mediated indels in EBE-Type I CsLOBP in transgenic Valencia Line #V5. The I CsLOBP sequence targeted by Cas9/sgRNA:CsLOBP2 was shown in blue, and the mutations were shown in purple. The protospacer-adjacent motif (PAM) was indicated in red. The PthA4 effector binding elements were highlighted by red rectangle. Among 50 sequenced colonies, 18 correspond to Type I CsLOBP, 19 to Type II CsLOBP and 13 to mutant Type I CsLOBP. (b) The chromatograms of EBE-Type I CsLOBP and its edited variants. The targeted sequence within the Type I CsLOBP was indicted by a black line, and the mutant sites were shown by arrows.

The three transgenic Valencia lines generated via mature tissue transformation and wild type Valencia plants were inoculated with Xcc at a concentration of 5×10^8^ CFU/ml. At 3 days post inoculation (DPI), typical canker symptoms were observed on both wild-type Valencia and all three transgenic lines (Fig. 5a). By 6 DPI, the symptoms became more severe in all plants (Fig. 5b). In contrast, XccΔpthA4:dCsLOB1.3 caused similar canker symptoms on WT, #V3 and #V5, but did not cause canker symptoms on #V2 at both time points (Fig. 5). This is consistent with that induction of one *CsLOB1* allele is sufficient to induce citrus canker symptoms (Jia et al., 2016). It is noteworthy that XccΔpthA4:dCsLOB1.3 carries the artificial dTALE dCsLOB1.3 that specifically activates Type I CsLOBP, but not Type II CsLOBP (Jia et al., 2016).

**Figure 5.**
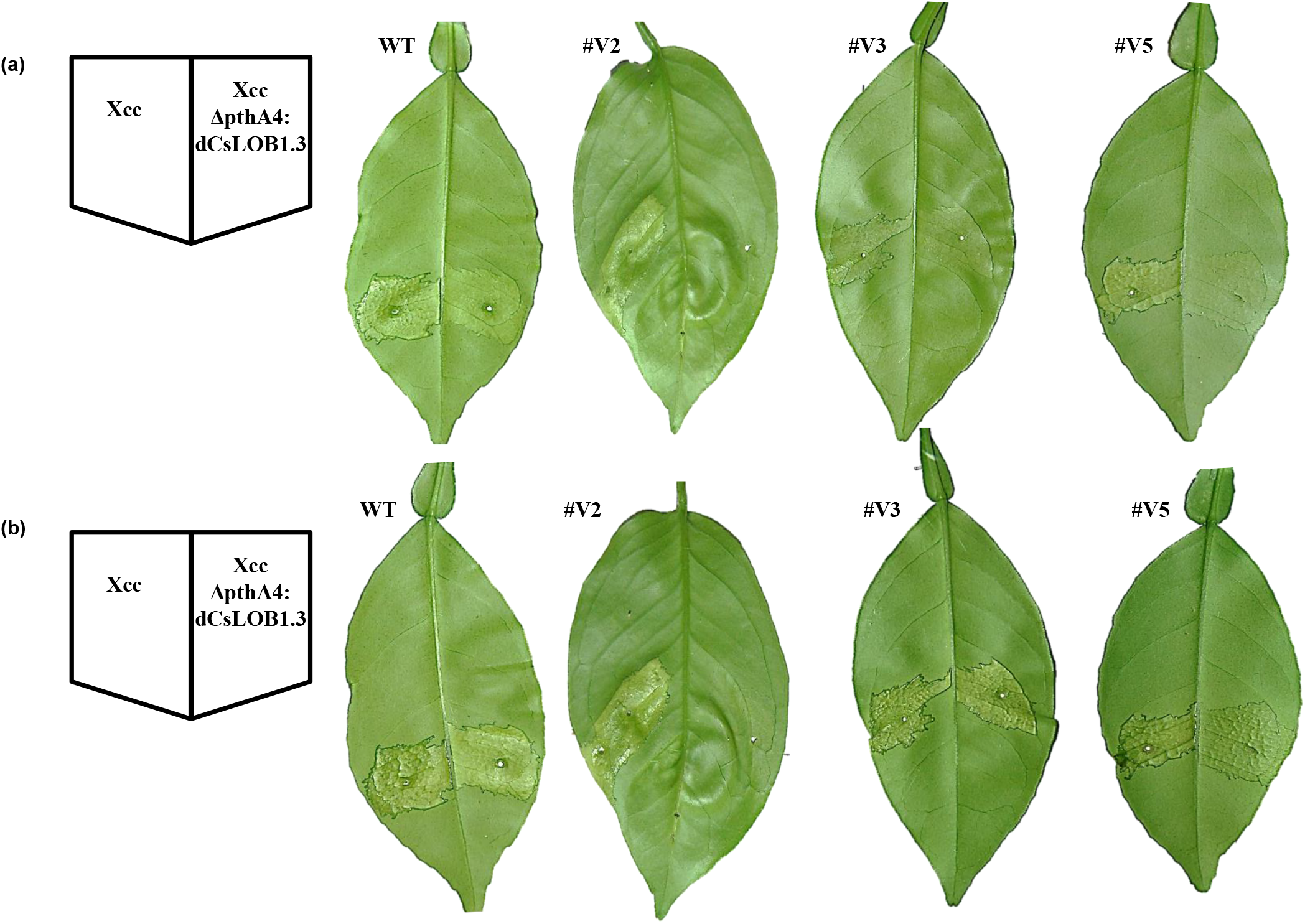
Differential canker-inducing responses of Valencia and EBE_PthA4_-TI CsLOBP edited lines to Xcc and Xcc306ΔpthA4:dCsLOB1.3. (a) At 3 days post inoculation (DPI) with Xcc, there were typical canker symptoms on wild type Valencia and transgenic lines. Wild-type Valencia served as the susceptible control. Following inoculation with Xcc306ΔpthA4:dCsLOB1.3, no canker symptoms were observed on line #V2 at 3 DPI, consistent with its 100% mutation rate in EBE_pthA4_-TI CsLOBP, whereas weak symptoms were detected on wild type Valencia, #V3 and #V5. Lines #V3 and #V5 exhibited lower mutation rates than #V2. (b) At 6 DPI with Xcc, severe canker symptoms developed on #V2, #V3, #V5, and wild type Valencia. However, at 6 DPI with Xcc306ΔpthA4:dCsLOB1.3, there was still no canker on line #V2, although canker symptoms became more evident on #V3, #V5, and wild-type Valencia. These results indicate that #V2 is resistant to Xcc306ΔpthA4:dCsLOB1.3.

Transgenic *LOB1* edited citrus plants regenerated from mature tissues grew normally and did not show obvious defects in growth and development. All three transgenic Valencia plants initiated flowering in 15 months whereas transgenic lines generated via embryogenic protoplasts and epicotyl transformation did not flower when kept for 3 years in the greenhouse conditions (Fig. 6). Consistently, mature transgenic sweet orange plants flowered in 14 months after establishment (Cervera et al., 1998).

**Figure 6.**
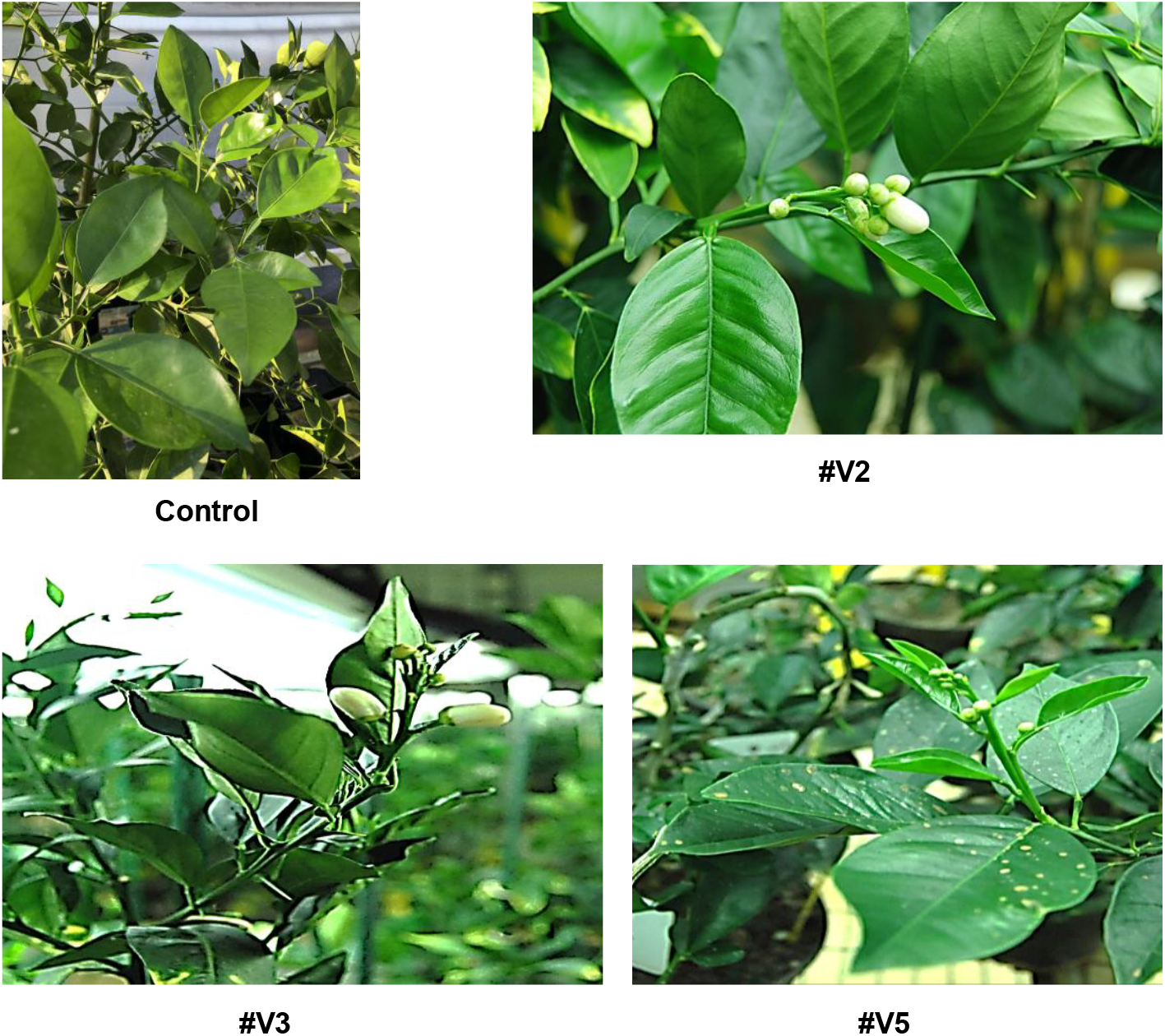
Early flowering of EBE_PthA4_-TI CsLOBP edited Valencia lines generated via mature tissue transformation. *LOB1* edited sweet orange generated via embryogenic protoplast transformation did not flower at 3 years under greenhouse conditions. Transgenic Valencia lines (#V2, #V3, and #V5) flowered within 15 months under greenhouse conditions.

## Discussion

In this study, we have shown for the first time that Cas9/sgRNA-mediated genome editing of citrus via mature tissue transformation achieves both high-efficacy genome editing and early flowering. While annual plants have a shorter juvenile phase, perennial trees have a long juvenile phase up to several decades (Pan et al., 2023). Among them, citrus species have a juvenile phase of 2-20 years with sweet orange around 5-10 years before they can flower and bear fruit when grown from seeds or other juvenile tissues (Gmitter et al., 2012). Such an obstacle has significantly delayed the evaluation of tree plants generated via biotechnical approaches and their eventual adoption for commercial production. Mature tissue transformation is a promising method to shorten juvenility of citrus. Cervera et al. first established the mature tissue transformation for citrus by transgenic expression of a *uidA* reporter in Pineapple sweet orange, which flowered 14 months after transformation (Cervera et al., 1998). Cervera et al. further conducted successful transformation of Clementine mandarin with a GFP reporter using mature tissues, which started to flower 6-9 months after grafting (Cervera et al., 2008). The plants flowered and set fruits during consecutive springs. Mature tissue transformation was also successfully used for transgenic expression in sweet orange cultivars Pera, Natal, Valencia, Hamlin, Florida EV1, Ruby Red grapefruit, citron (*C. medica*), rootstock varieties US942 and Kuharske (Almeida et al., 2003; Canton et al., 2024; Canton et al., 2022; Marutani-Hert et al., 2012; Wu et al., 2015). Importantly, mature tissue transformation was used to generate transgenic sweet orange expressing *Cecropin* B and *Shiva* A and 11 of the 40 transgenic lines showed enhanced disease resistance to citrus canker (He et al., 2011). Additionally, transgenic expression of RNA interference (RNAi) constructs targeting specific CTV sequences was conducted to generate CTV resistant citrus plant via mature tissue transformation (Soler et al., 2019). However, none of these studies conducted CRISPR genome editing through mature tissue transformation.

CRISPR genome editing has demonstrated tremendous potential in genetic improvement of citrus. It was successfully used to improve citrus resistance against canker by editing the canker susceptibility gene *LOB1* (Huang Wang 2021 (Huang et al., 2021; Jia et al., 2021; Jia et al., 2017b; Peng et al., 2017); Huang Wang 2022; Jia Wang 2021; Jia Wang 2017, Peng 2017) and *DMR6*, which encodes a salicylic acid 5-hydroxylase (Parajuli et al., 2022). It has also been used to improve citrus resistance/tolerance to HLB by targeting *NPR3* (Tiwari et al., 2024) and *EDS1* (Huang et al., 2025). Furthermore, it has also been used for non-transgenic genome editing of citrus (Alquézar et al., 2022; Huang et al., 2023; Huang et al., 2022; Rocha et al., 2025; Su et al., 2023). Despite the significant progress of CRISPR genome editing in citrus genetic improvement, previous citrus genome editing studies have relied exclusively on immature explants, such as epicotyls, callus, and protoplasts (Prado et al., 2024). In this study, we demonstrated that Cas9/sgRNA-mediated genome editing of citrus via mature tissue transformation can achieve both high-efficacy genome editing and early flowering, thus accelerating the evaluation and deployment of genome edited citrus. Among the three genome edited citrus lines, one achieved 100% genome editing efficacy of the target sequence EBE_pthA4_-TI CsLOBP. No off-target mutations were detected for EBE_pthA4_-TII CsLOBP, which has only one nucleotide difference from EBEpthA4-TI CsLOBP, demonstrating high fidelity of the Cas9/gRNA mediated genome editing of citrus. We, however, did not acquire biallelic/homozygous mutations that were successfully generated using either epicotyl or embryogenic protoplasts (Huang et al., 2020; Jia et al., 2021; Su et al., 2023). In addition, the plants generated in this study were transgenic. It remains to be explored whether mature tissue transformation can be used to generate non-transgenic genome-edited citrus for easing the deregulation hurdles and better public acceptance (Huang et al., 2023; Su et al., 2023).

## Acknowledgements

We thank Wang lab members for constructive suggestions and insightful discussions. This work is supported by the Emergency Citrus Disease Research and Extension program, project award no. 2022-70029-38471, 2021-67013-34588, 2025-70029-44030 and 2023-70029-41280 from the U.S. Department of Agriculture’s National Institute of Food and Agriculture, J. R. Graves Endowment, Florida Citrus Initiative Program, Citrus Research and Development Foundation, and Hatch project [FLA-CRC-005979] (N. Wang).

## Data Availability Statement

The data that support the findings of this study are available in the Supporting Information of this article.

