## Supplementary Figures for "Cas9/sgRNA-mediated genome editing of citrus via mature tissue transformation enables both high-efficacy genome editing and early flowering"

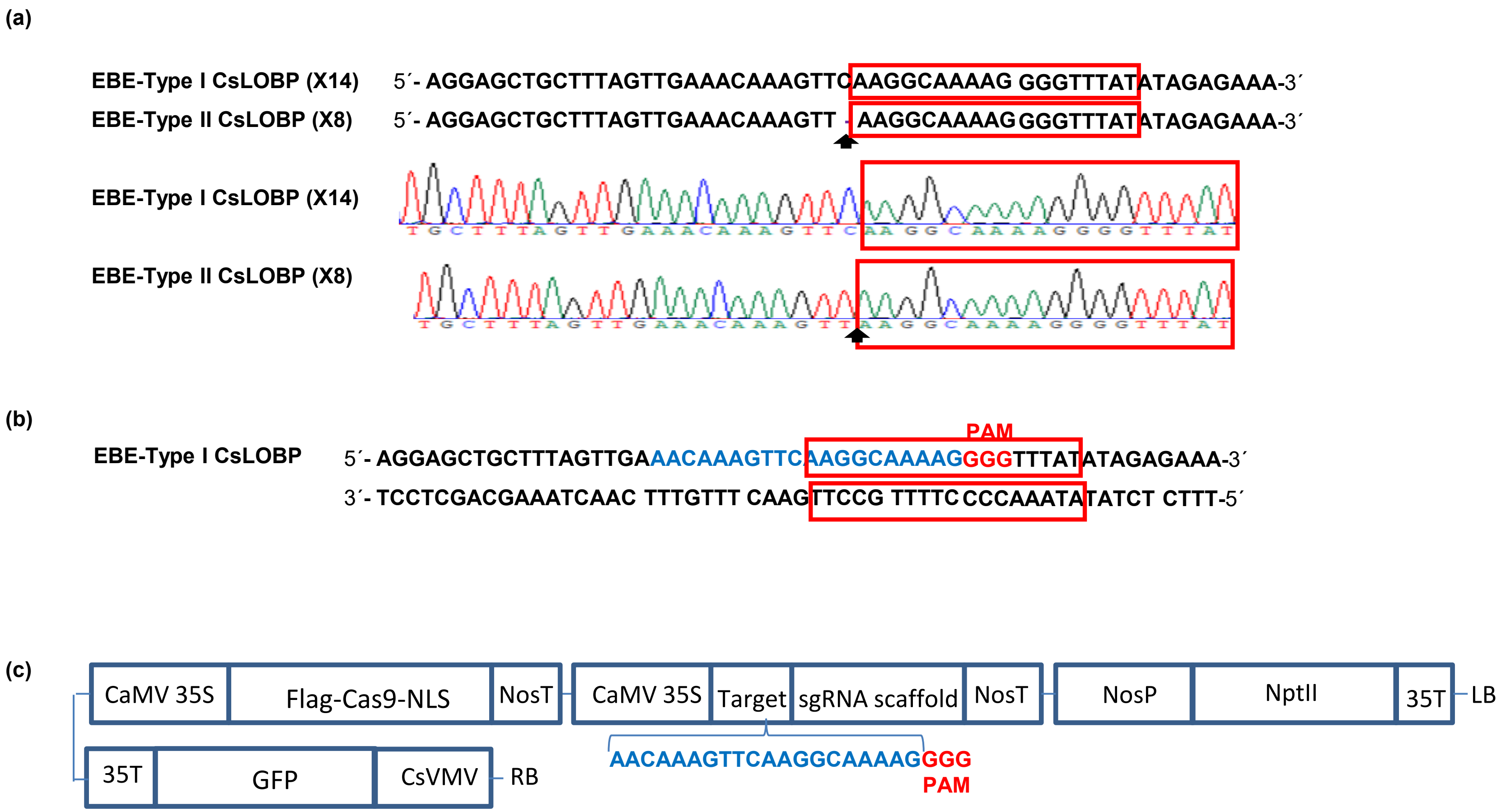


**Supplementary Figure 1. Valencia CsLOBP region and binary vector GFP-p1380N-Cas9/sgRNA:CsLOBP2.** (a) Two types of CsLOBP (Type I and Type II) identified in Valencia and their representative chromatograms. The 1-bp difference between the two types is indicated by arrows, and the PthA4 effector binding elements were highlighted with red rectangles. Among 22 sequenced colonies, 14 correspond to Type I CsLOBP, and 8 correspond to Type II CsLOBP. (b) A single-guide RNA (sgRNA:CsLOBP2) was designed to target the $EBE_{pthA4}$-TI CsLOBP. The sgRNA:CsLOBP2 target site was highlighted in blue. The protospacer-adjacent motif (PAM) was indicated in red. (c) Schematic representation of the GFP-p1380N-Cas9/sgRNA:CsLOBP2 construct. LB and RB indicate the left and right borders of the T-DNA region. CsVMV, cassava vein mosaic virus promoter; GFP, green fluorescent protein; CaMV 35S and 35T, cauliflower mosaic virus 35S promoter and terminator; NosP and NosT, nopaline synthase promoter and terminator; Flag-Cas9-NLS, Cas9 endonuclease fused with an N-terminal Flag tag and a C-terminal nuclear localization signal; target, the 20 nucleotides of $EBE_{pthA4}$-TI CsLOBP highlighted by blue, was located upstream of PAM; sgRNA scaffold, a synthetic single-guide RNA composed of a fusion of CRISPR RNA and *trans*-activating CRISPR RNA; NptII, neomycin phosphotransferase II.

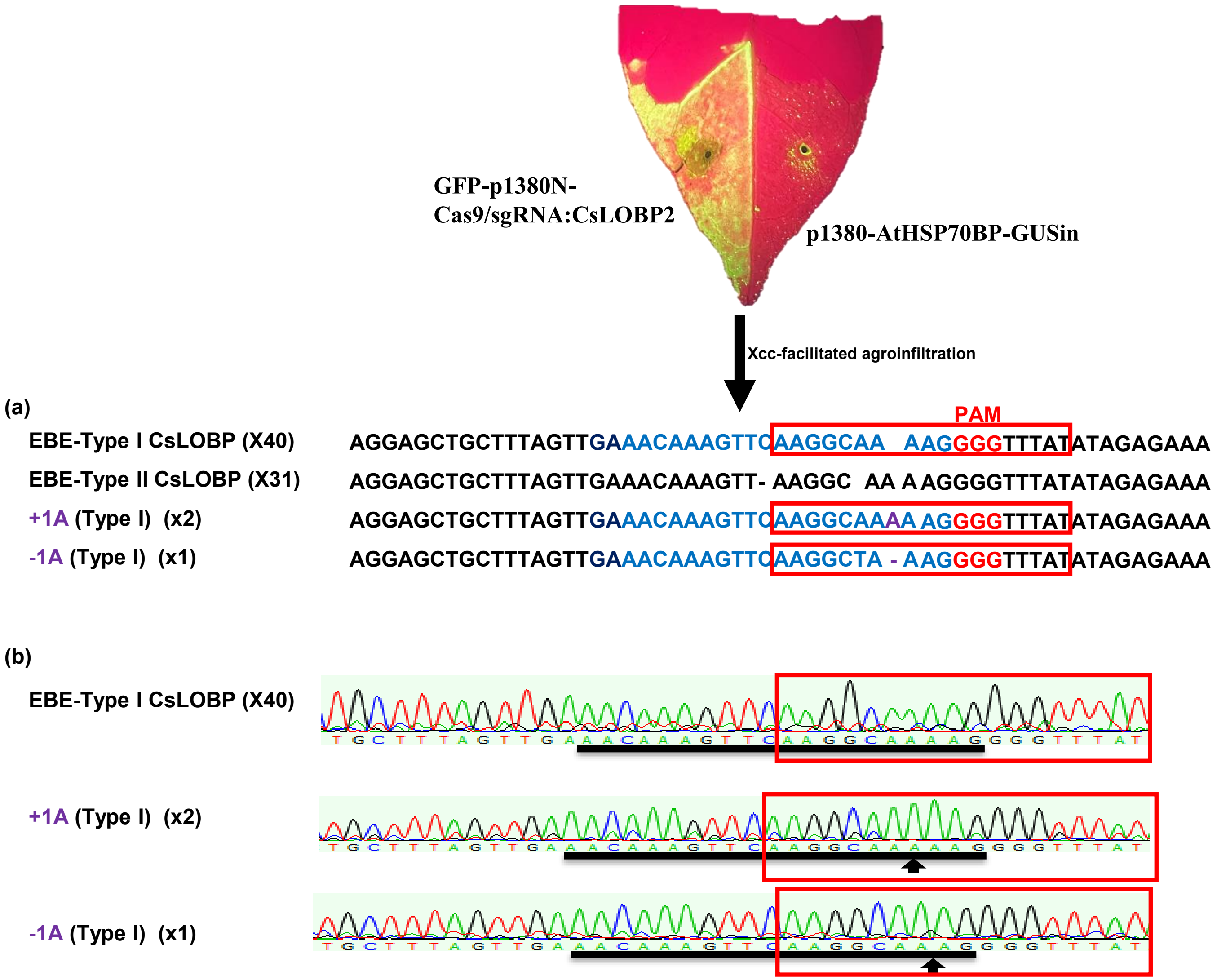


**Supplementary Figure 2. Analysis of Cas9/sgRNA:CsLOBP2-mediated indels in Valencia leaves using Xcc-facilitated agroinfiltration.** (a) Targeted mutations induced by Cas9/sgRNA:CsLOBP2 in EBE$_{pthA4}$-TI CsLOBP in Valencia. The sgRNA:CsLOBP2 target sequence in Type I CsLOBP was highlighted in blue, and induced mutations were indicated in purple. The protospacer-adjacent motif (PAM) was indicated in red. The PthA4 effector binding elements were marked with red rectangles. Among 74 sequenced colonies, 40 correspond to Type I CsLOBP, 31 to Type II CsLOBP, and 3 to mutated Type I CsLOBP. These results indicated that Cas9/sgRNA:CsLOBP2 specifically targeted Type I CsLOBP. (b) Representative chromatograms of Type I CsLOBP and its edited variants. The mutation site was indicated by an arrow, and the sgRNA target region within Type I CsLOBP was denoted by a black line.
